# Hypocotyl Development in Arabidopsis and other Brassicaceae Displays Evidence of Photoperiodic Memory

**DOI:** 10.1101/2024.05.13.593876

**Authors:** James Ronald, Sarah C.L Lock, Will Claydon, Zihao Zhu, Kayla McCarthy, Elizabeth Pendlington, Ethan J. Redmond, Gina Y.W. Vong, Sanoj P. Stanislas, Seth J. Davis, Marcel Quint, Daphne Ezer

## Abstract

Sensing and responding to photoperiod changes is essential for plants to adapt to seasonal progression. Most of our understanding of how plants sense photoperiodic changes is through studies on flowering time. However, other aspects of plant development are regulated by the photoperiod, including hypocotyl elongation. Unlike flowering, hypocotyl elongation displays a greater plasticity to changes in the photoperiod with increases in daylength causing greater inhibition of growth until a threshold is met. Previous studies have only looked at hypocotyl development in the context of a stationary photoperiod. It is unknown if changes in the photoperiod during development influence hypocotyl elongation. Here, we developed a physiological assay to investigate this question. We have discovered that hypocotyl elongation is influenced by a memory of past photoperiod exposure in Arabidopsis and Brassicaceae cultivars used for microgreen agriculture. Photoperiodic memory persisted for multiple days, although it weakened over time, and the strength of the memory was dependent on the genetic background. We identified that phyB and ELF3, key regulators of hypocotyl development, were required for photoperiodic memory. Finally, we identified that the circadian clock is unlikely to function as a repository for photoperiodic memory as circadian rhythms quickly re-aligned with the new photoperiod. In summary, our work highlights for the first-time evidence of a photoperiodic memory that can control plant development.

## Introduction

The daily rotation of the Earth generates cycles of light and dark. The magnitude change in these daily cycles across the year is influenced by the distance to the equator, with organisms in latitudes further from the equator experiencing greater extremes in photic and thermal change than those at the equator. Many organisms have adopted a suite of mechanisms to perceive and respond to these seasonal changes, particularly changes in the photoperiod. Adaptation to seasonal changes is critical for organisms to maximize their fitness and productivity.

In plants, photoperiodic responses have been best studied in the context of the transition from vegetative to floral growth. This has led to the establishment of the coincidence model to describe how the photoperiod controls the transition to flowering (Song et al., 2015). However, other aspects of plant development are also controlled by the photoperiod. This includes a recently reported metabolic photoperiod perception pathway that regulates growth under long-day (LD) photoperiods (Wang et al., 2024). Additionally, the development of the hypocotyl is regulated by the photoperiod (Krahmer and Fankhauser, 2023). Similar to photoperiodic regulation of flowering, a coincidence model has been proposed to explain how hypocotyl elongation is controlled by the photoperiod (Nozue et al., 2007; Niwa, Yamashino and Mizuno, 2009). Within the hypocotyl coincidence model, PHYTOCHROME INTERACTING FACTOR4 (PIF4) and PIF5 function as positive regulators of hypocotyl elongation (Nozue et al., 2007). The timing of *PIF4/5* expression is regulated by the circadian clock in a photoperiod-dependent manner (Niwa, Yamashino and Mizuno, 2009). Maximal hypocotyl elongation occurs under short-day (SD) photoperiods as *PIF4/5* expression coincides with the dark, avoiding the repressive effect of light-activated photoreceptors (Leivar and Monte, 2014), and day and dusk-phased circadian proteins on PIF4/5 transcriptional activity (Nieto et al., 2015; Martín et al., 2018; Nohales et al., 2019).

Although flowering and hypocotyl development share a similar model to describe how the photoperiod is perceived and then acted upon, the effect of the photoperiod on the transition to flowering and hypocotyl development has fundamental differences. Photoperiodic-dependent induction of flowering is a binary process, whereby once the photoperiod meets the threshold flowering is initiated (Song et al., 2015). In contrast, hypocotyl development is more plastic to the photoperiod. Increasing the length of the photoperiod causes greater inhibition of hypocotyl elongation until a threshold is met where further increases in light duration has no additional inhibitory effect (Niwa, Yamashino and Mizuno, 2009; Anwer et al., 2020). The current photoperiodic model of hypocotyl elongation describes development exclusively in the context of a stable photoperiod (Nozue et al., 2007; Niwa, Yamashino and Mizuno, 2009). This model has not been tested under conditions where the photoperiod duration changes during development. Moreover, it is unknown if photoperiod transitions during development can influence hypocotyl development, a so-called “memory” of past-events. Such memory may have important implications in microgreen agriculture where hypocotyls function as the commercial product. Here, balancing the photoperiod in microgreen agriculture is important to maximize the nutritional value but also minimizing the energy requirements and associated costs from growth under LD photoperiods (Bhaswant et al., 2023; Filatov and Olonin, 2023; Hernández-Adasme, Palma-Dias and Escalona, 2023).

In this manuscript, we investigated whether plants retain memory of prior photoperiod exposure using hypocotyl development as a physiological system. We have designed a photoperiod shift assay to assess how prior photoperiod exposure influences hypocotyl elongation in *Arabidopsis thaliana* (Arabidopsis hereafter) seedlings and germinating seeds. We have found that Arabidopsis displays evidence of a photoperiodic memory. The strength of the memory was dependent on the direction of the photoperiod shift and the developmental age of the Arabidopsis plants when they were initially entrained to the first photoperiod. We have identified key roles for phytochromeB (phyB) and EARLY FLOWERING3 (ELF3) in regulating photoperiodic memory response. We also investigated whether circadian rhythms could encode information on prior photoperiodic exposure. However, circadian rhythms quickly adapted to changes in photoperiod, suggesting that other mechanisms are required for encoding historical photoperiod information. Finally, we identified that several vertically farmed Brassicaceae (kale and radish) have stronger photoperiodic memory than Arabidopsis. Ultimately, hypocotyl memory could have important applications in developing energy-efficient growth regimes for microgreen commercial applications.

## Methods

### Plant lines and husbandry

The following *Arabidopsis thaliana* ecotypes were used in this work: Bayreuth-0 (Bay-0), Cape Verde Island (Cvi), Sapporo (Sap), Shakdara (Sha), Tanzania-1 (Tnz-1) and Wassilewskija-2 (Ws-2). All ecotypes harbored the *CCR2::LUC* reporter gene. Bay-0 *CCR2::LUC*, Cvi *CCR2::LUC*, Sha *CCR2::LUC* and Ws-2 *CCR2::LUC* have been described previously (Doyle et al., 2002; Boikoglou et al., 2011; Anwer et al., 2014). For the Sap and Tnz *CCR2::LUC* lines, the respective parent backgrounds were transformed with the *CCR2::LUC* reporter by floral dip (Davis et al., 2009). A luminesce transgenic plant was then identified and backcrossed to the respective parental line. This backcrossed transgenic was then self-fertilized before a homozygous, single transgene segregant was identified and bulked in the following generation. The following Arabidopsis mutants in the Ws-2 background were also used: *elf3-4 CCR2::LUC*, *phyB-10 CCR2::LUC*, *elf3-4*/*phyB-10 CCR2::LUC*. These mutants have been previously described in earlier work (Reed et al., 1993; Hicks, Albertson and Wagner, 2001; Ronald et al., 2022). Prior to experiments, all lines were propagated on soil in environmentally controlled growth chambers with a LD photoperiod (16/8 light/dark) at a light intensity of 100 µmol/m^-2^/s^-1^ and a set temperature of 22°C.

### Hypocotyl measurements

For the collection of data presented in figure 1, figure 2 and supplementary figure 1-5, all seeds were surface sterilized before being plated onto 1x Murashige and Skoog (MS) media with 1.5% w/v phytoagar (Duchefa), 1% w/v sucrose (Sigma) and 0.5 g/L MES (Sigma). Seeds were then stratified for 48-96 hours. After stratification, germination was induced by exposure to 75 µmol/m^-2^/s^-1^ of white light for 24 hours and a constant temperature of 19.5°C. After 24 hours, germinating seeds were provided photoperiod information. They were either grown under SD (6/18 light/dark) or LD photoperiods with a constant temperature of 19.5°C and a light intensity of 75 µmol/m^-2^/s^-1^. After 48 hours, the photoperiod shifts were initiated. Seedlings were either kept under the initial photoperiod as a control or transferred to the reciprocal photoperiod. All photoperiod shifts occurred at ZT6, before lights off for the SD growth conditions. Measurements were taken on day 5 (timepoint 1) or day 7 (timepoint 2) at ZT6. All experiments were repeated twice with a similar result observed across the two experimental repeats. The presented data is a combination of the two experimental repeats, with each individual experiment including at least fifteen seedlings. Hypocotyl images were measured using ImageJ (Schneider et al. 2012). For figures 1 and 2, hypocotyl measurements were scaled so that the median LD hypocotyl length was 0 and the median SD hypocotyl length was 1, using the formula: *Scaled length = (length – median length LD)/(median length SD – median length LD).* Adjusted p-values were calculated using TukeyHSD tests. We have included all statistical and raw hypocotyl measurements as a supplementary dataset (Supplementary File).

**Figure 1.**
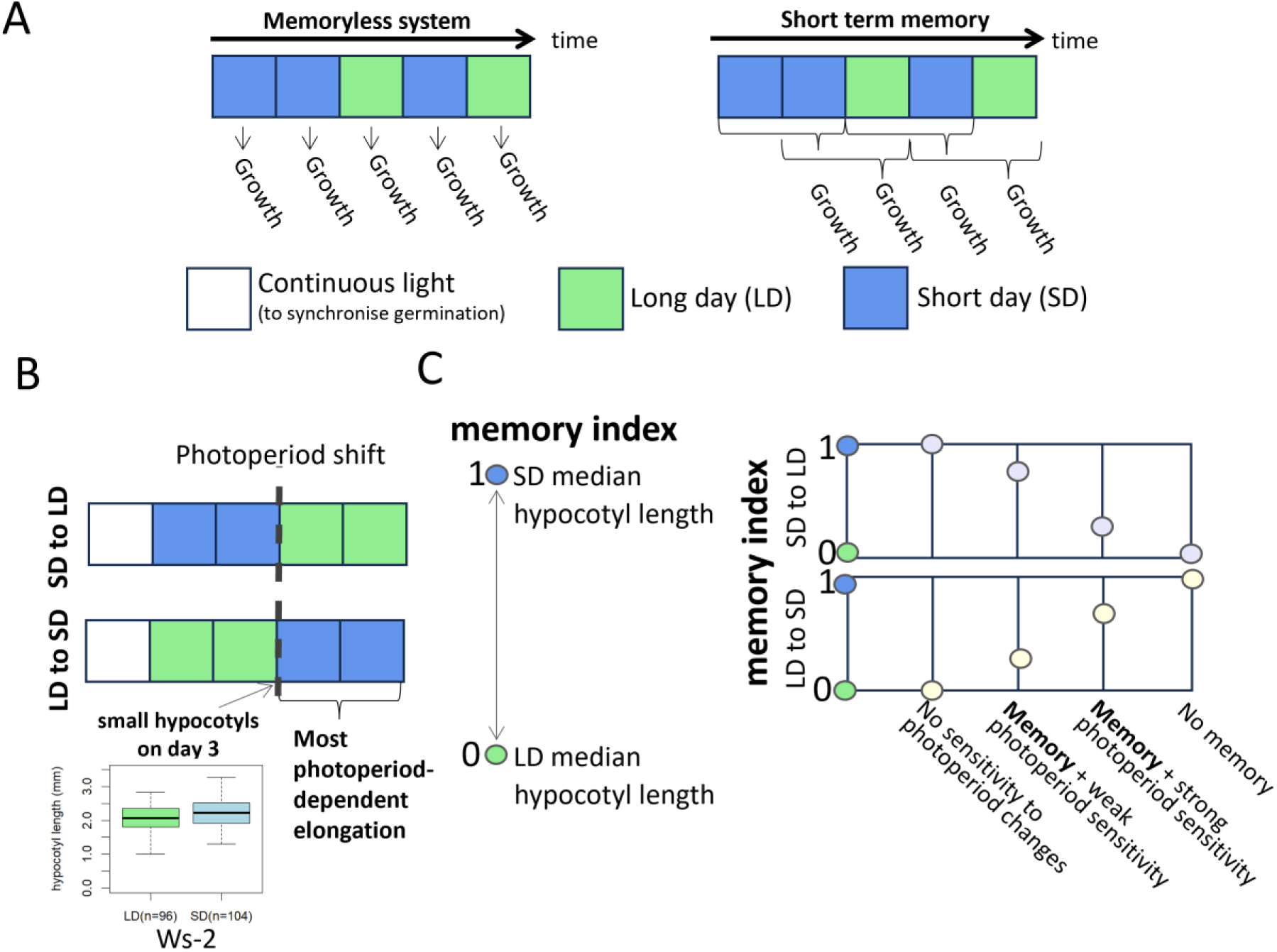
Designing an assay to test for photoperiodic memory. (A) In the context of this manuscript, we define a ’memory-less system’ as one whose growth only depends on the current photoperiod. Conversely, ’memory’ refers to growth that also depends on prior environmental conditions. (B) Hypocotyl growth in Arabidopsis seedlings occurs predominantly during the period after the photoperiod shift has occurred. (C) A hypocotyl memory index was used to compare photoperiodic memory and the sensitivity to photoperiodic shifts between ecotypes and mutants that have large magnitudes differences in hypocotyl elongation. Hypocotyl lengths were scaled so that the median hypocotyl length at LD (16/8hr light/dark) is 0 and the median hypocotyl length at SD (8hr/16hr light/dark) is 1.

**Figure 2.**
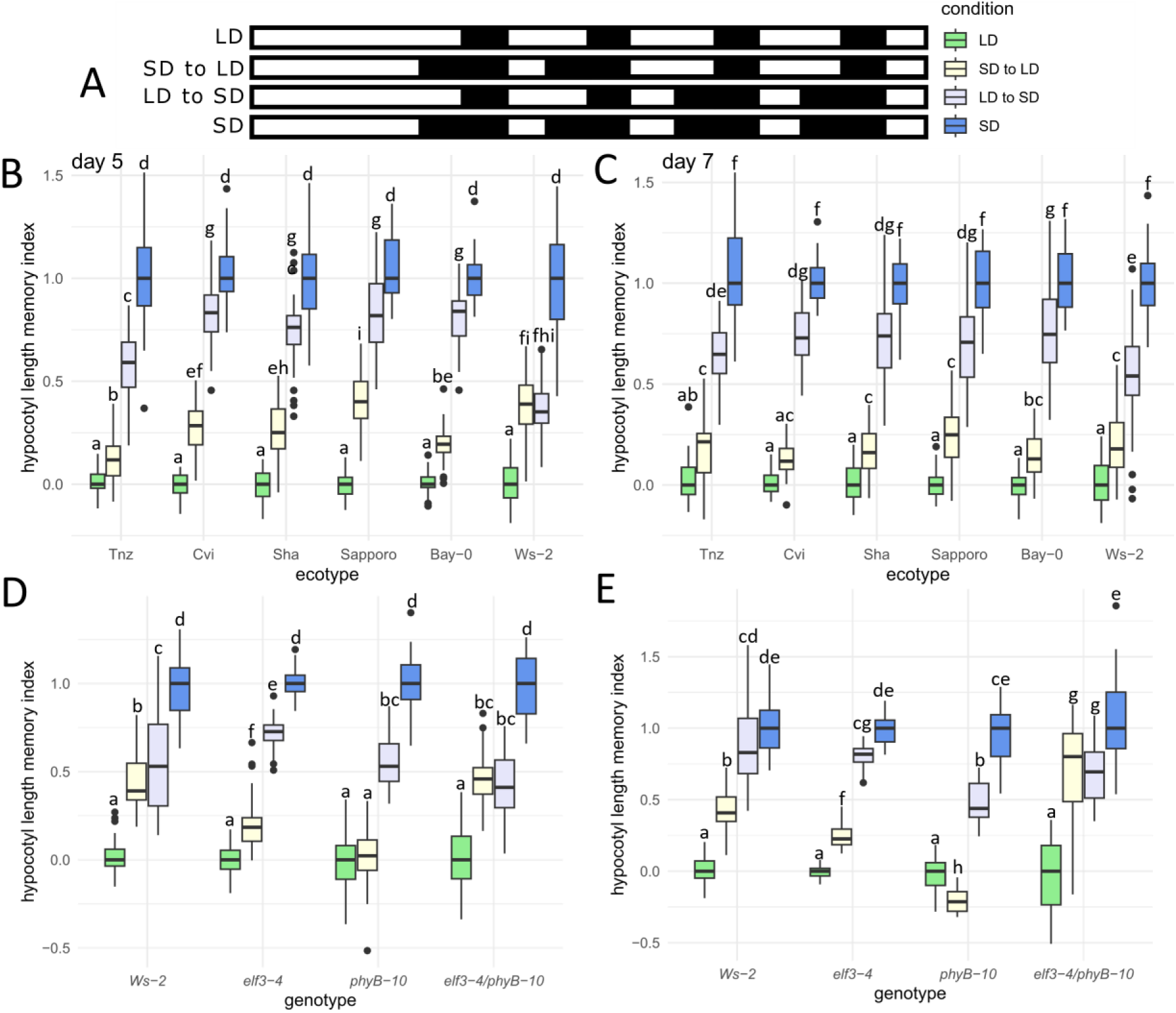
Hypocotyl development in Arabidopsis displays evidence of photoperiodic memory. (A) Diagram explaining the four different photoperiodic growth conditions, the day phase is white, and the night phase is black. Different Arabidopsis wild-type ecotypes were either measured (B) two or (C) four days after the respective photoperiod shift or were kept under the same photoperiod for the duration of the experiment. Photoperiodic memory response in *elf3-4*, *phyB-10* and *elf3-4/phyB-10* mutants after either (D) two days or (E) four days following the photoperiod shift. Letters signify a significant adjusted p-value of p < 0.01, determined by TukeyHSD.

### Luciferase assay

Arabidopsis Ws-2 *CCR2::LUC* WT seeds were surface sterilized and plated on 1x MS medium containing 3% w/v sucrose (Sigma), 1.5% w/v phytoagar (Duchefa), 0.5 g/L MES (Sigma) and 15 µg/ml Hygromycin-B (Duchefa). After 5 days of stratification at 4°C, seeds were transferred to the specified photoperiod for 6 days. For all photoperiods, a constant temperature of 20°C was used and a light intensity of between 80 - 100 µmol/m^-2^/s^-1^. On day 6, seedlings were transferred into a 96 well plate containing the same media as described above but additionally 15 µl of 1 mM Luciferin. Plates were then placed back in the growth cabinet until they were then transferred to the TOPCOUNT (TC) at dawn on day 7. Plates in the TC were grown under the stated photoperiod with red plus blue light (2.4-3.6 µmol/m^-2^/s^-1^ red light and 0.8-1.4 µmol/m^-2^/s^-1^ blue light) and a constant temperature of 20°C for a further 4 days. For samples undergoing a photoperiod shift, this shift occurred on day 11, exactly 96 hours after being placed in the TC. The fda package in R was used to smooth curves using B-splines and perform functional principal component analysis.

### Continuous monitoring of hypocotyl elongation

Sterilized Arabidopsis seeds were stratified in darkness at 4°C for 72 hours. To avoid contact of seedlings with the media surface during hypocotyl elongation, two strips were cut and removed from the cultivation medium. Solid *A. thaliana* solution (ATS) nutrient medium (Lincoln, Britton and Estelle, 1990) with 1% w/v sucrose was used as the cultivation media for these experiments. Seeds were sown in small indentations on the surface of the cut to prevent excessive movement as described previously (Janitza et al., 2024). Plates were placed vertically at 21°C and constant light (LL, 40 μmol m^-2^s^-1^) for one day to allow uniform germination of seeds, before being transferred to the initial photoperiod: either LD or SD conditions. After two days, plates were transferred to the alternative photoperiod (SD or LD, dawn-aligned), or were kept under the initial photoperiod. The analysis of hypocotyl elongation then started 72 hours after sowing (ZT0). Seedlings were illuminated by two infrared light emitting diodes and were imaged hourly by a Panasonic G5 camera modified with a long pass filter (830 nm). To detect minor changes between time points, image stacks were imported as “Image Sequence” in ImageJ. Each seedling (n≳6) was analyzed separately, and the hypocotyl length was measured using the “Segmented Line” tool, by slightly adjusting the control nodes between time points.

### Hypocotyl Elongation in Microgreens

Microgreens (Supplementary Table 2) seeds were sown into trays containing a compost mix of equal parts F2+S (Levington), coir and vermiculite supplemented with 0.7 g/L osmocote fertilizer. Seeds were sown approximately 1 cm below the surface of the soil. Microgreens were then grown under either a LD or SD photoperiod with a constant temperature of 19°C, 60% humidity and 70 µmol/m^-2^/s^-1^ of light. Trays were inspected daily to determine when individual hypocotyls had emerged from the soil. Once 50% of the respective microgreen cultivar had a visible hypocotyl, seedlings were grown for three days under the initial photoperiod. This period was determined as the initial photoperiod exposure. After this period, the microgreens were then either switched to the reciprocal photoperiod or kept under the same photoperiod as a control for a further 3 days. Trays were switched at ZT5 to ensure that most of the plants that underwent photoperiod shifts had equal exposure to each photoperiod. Each experiment was carried out three times with 40 individual plants imaged per variety per experiment. To image the microgreen hypocotyls, leaves were firstly dissected before the hypocotyls were removed from the soil and laid flat in an Epson scanner. High-resolution scans were then taken. Images were imported into ImageJ for measurement. All imaging took place at ZT5.

## Results

### Designing a photoperiodic memory experimental assay

To understand whether hypocotyl development in Arabidopsis is influenced by prior exposure to a given different photoperiod (Figure 1A) we designed the following assay. Stratified seeds were firstly germinated under constant white light for 24 hours to synchronize germination before seedlings were then provided two days of photoperiodic information, either SD or LD. On day 3, seedlings were then either transferred to the reciprocal photoperiod or kept under the existing photoperiod. Hypocotyl measurements were then taken on day 5 and day 7 to understand the immediate and long-term impact of the shift, respectively. We transferred seedlings on day 3 as hypocotyl development in de-etiolated Arabidopsis seedlings primarily occurs between three- and four-days post-germination (Gendreau et al., 1997). Supporting this, we found minimal difference in median hypocotyl length of Ws-2 seedlings grown under SD and LD on day 3 (Figure 1B). To compare the relative amount of photoperiodic memory between Arabidopsis ecotypes and mutants with different hypocotyl lengths, we implemented a photoperiodic memory index (PMI, Figure 1C). The PMI scales the hypocotyl length relative to the median hypocotyl length of growth under LD and SD conditions. Thus, seedlings with a PMI closer to 1 have a hypocotyl length similar to those grown under constant SD, while a score closer to 0 indicates development similar to that of constant LD.

### Arabidopsis seedlings display photoperiodic memory

We firstly analyzed the photoperiodic memory of wild-type Arabidopsis Ws-2 seedlings. As discussed above, Ws-2 seedlings were exposed to the photoperiodic shift on day 3 (Figure 2A). By day 5, hypocotyl development in the Ws-2 seedlings that underwent either photoperiod shift had a PMI of ∼0.45 (Figure 2B, Supplementary Figure 1), indicating that exposure to the initial photoperiod does influence hypocotyl development. By day 7, hypocotyl development in the seedlings that underwent the photoperiod shift had diverged from each other and were more closely resembling those under constant conditions (Figure 2C). However, for neither photoperiod shift did hypocotyl elongation mirror that of seedlings grown under a constant photoperiod (Figure 2C). Together, these results suggest that hypocotyl development displays evidence of a lasting photoperiodic memory.

Next, we investigated whether other Arabidopsis ecotypes also displayed photoperiodic memory and the heterogeneity in response between these ecotypes. Ecotypes were picked to cover a range of latitudes from the equator to northerly latitudes (Supplementary Table 1). For all ecotypes, hypocotyl elongation again displayed evidence of photoperiodic memory and this memory continued to persist on day 7 (Figure 2B-C, Supplementary Figure 1). However, for the Sha, Cvi and Bay-0 ecotypes the memory of the initial photoperiod was weaker than Ws-2 for both photoperiod shifts. We also observed that certain ecotypes displayed stronger memory of a particular photoperiod shift. For example, Tnz had a stronger memory of the LD-SD than SD-LD transitions on day 5, while Sap displayed the opposite behavior (Figure 2B). However, neither effect persisted after an additional two days in the final photoperiod (Figure 2C). Importantly, for all ecotypes tested except Ws-2, there was a difference in hypocotyl length between the different photoperiod shifts on day 5 even though the seedlings were exposed to the same amount of total light (Supplementary Figure 2-3). Therefore, it is the order of the photoperiod exposure that is important in determining hypocotyl elongation.

### phyB and ELF3 regulate the rate that Ws-2 responds to changes in day length

To understand the genetic pathways underpinning the response to the photoperiodic shifts, we characterized different mutants in the Ws-2 ecotype background (wild type, WT). We firstly focused on phyB and ELF3 as they are key regulators of hypocotyl elongation (Reed et al., 1993; Zagotta et al., 1996). On day 5, the *elf3-4* mutant (*elf3* hereafter) had reduced photoperiodic memory of both photoperiod shifts relative to WT (Figure 2D, Supplementary Figure 4-5). On day 7, hypocotyl elongation in the *elf3* mutant remained more sensitive to the SD-LD shift but had a similar response to WT for the LD-SD shift (Figure 2E). For the *phyB-10* (*phyB* hereafter) mutant, seedlings phenocopied the WT response to the LD-SD on day 5 but then displayed reduced growth by day 7 (Figure 2D-E). Under the SD-LD shift, *phyB* mutants had a similar PMI to those kept under constant LD (Figure 2D), indicating a weak memory of the initial SD photoperiod. This phenotype became further enhanced by day 7, with *phyB* mutants moved from SD-LD now shorter than those kept under continuous LD (Figure 2E). We also analyzed the *elf3/phyB* double mutant phenotype to understand the genetic relationship between ELF3 and phyB. For the LD-SD shift, the *elf3/phyB* mutant phenocopied the WT response at both timepoints (Figure 2D-E). Similarly, for the SD-LD shift the *elf3/phyB* mutant had a similar response to WT at the first timepoint (Figure 2D). However, at the second time point the *elf3/phyB* mutant had reduced photoperiodic memory than WT (Figure 2E). Together, our results highlight a complex requirement of phyB and ELF3 in regulating photoperiodic memory: ELF3 is necessary to regulate a plant’s initial sensitivity to changes in the photoperiod, while phyB prevents over-compensation for the SD-LD shift and sensitizes plants to the shift to SD.

### The impact of photoperiodic memory is dependent on developmental age

Hypocotyl elongation is dependent on the accumulation of PIFs at specific times of day. Maximal hypocotyl elongation occurs under SD photoperiods as PIF accumulation coincides with the late evening, where PIF activity displays minimal inhibition (Niwa, Yamashino and Mizuno, 2009; Leivar and Monte, 2014). This led us to investigate how the dynamics of hypocotyl elongation changes across time in seedlings that underwent the photoperiod shift. For this experiment, seeds were stratified in water rather than on the surface of the agar-media for 3 days. This small change in the protocol delayed the onset of hypocotyl emergence until day 3, the point at which the photoperiod shift initiates. Thus, the perception of the initial photoperiod under these conditions occurred exclusively in the germinating seed. This allowed us to subsequently investigate how photoperiodic perception in seeds influences seedling development alongside the effects of the photoperiod shifts on the timing of hypocotyl elongation.

Intriguingly, WT seedlings whose germinating seeds were exposed to LD, but then shifted to SD had extremely rapid hypocotyl elongation, compared to all other conditions (Figure 3A). The timing of growth in seeds that were shifted from LD-SD was consistent with those grown under constant SD conditions (Figure 3B). However, the maximal growth rate in the seeds that were shifted from LD-SD was much higher on the first night and remained elevated on the second night. Thus, the shift from LD-SD caused a temporary acceleration of growth. For the SD-LD shift, WT displayed a less pronounced memory of the initial SD exposure, with overall hypocotyl elongation (Figure 3A) and the timing of growth (Figure 3B) closely mirroring seedlings that were only grown under constant LD. In summary, WT seeds display photoperiodic memory of prior LD exposure, and this results in sustained levels of increased hypocotyl elongation when shifted to SD.

**Figure 3.**
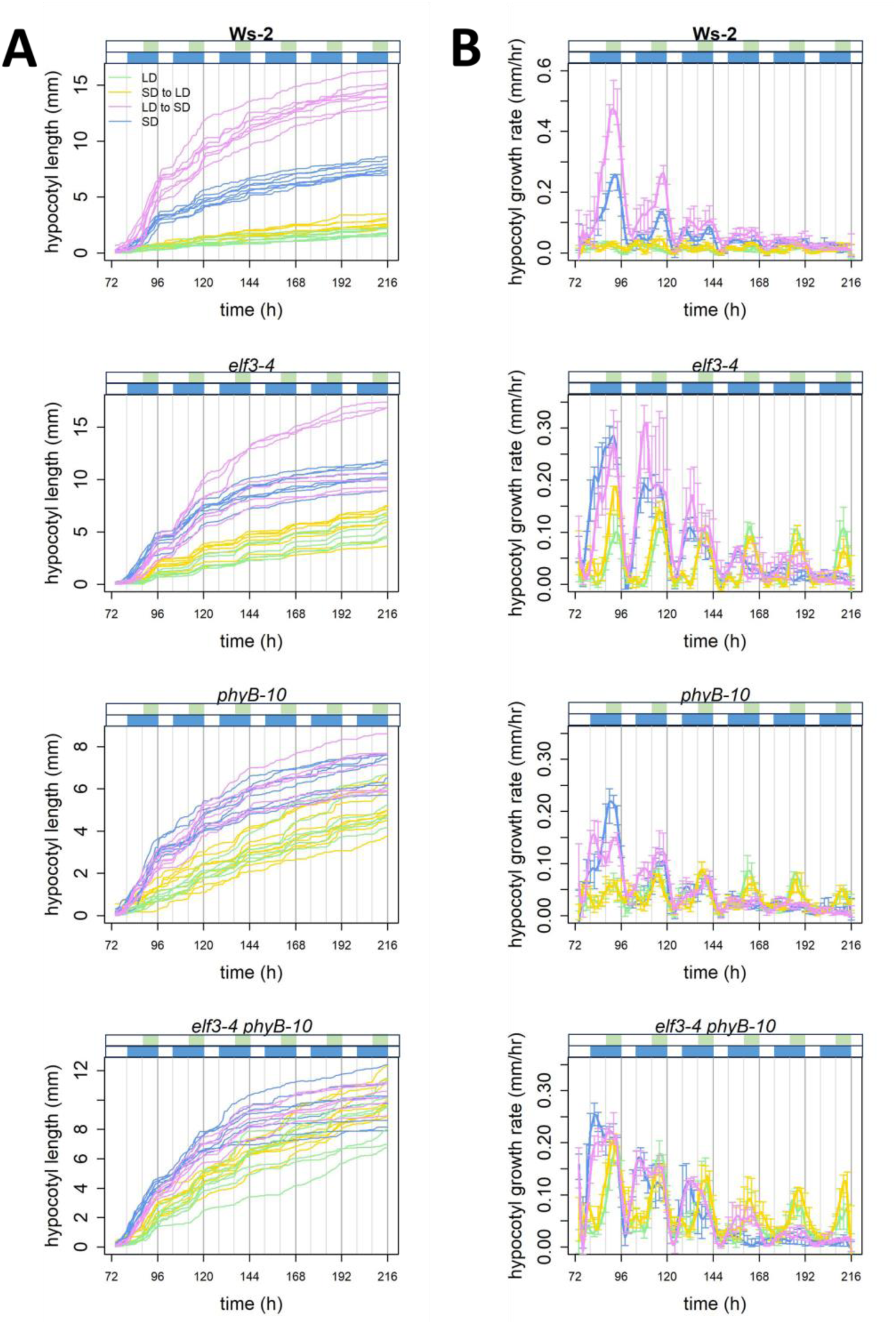
Photoperiodic memory response is dependent on the developmental age of the plant. (A) Hypocotyl length in Arabidopsis Ws-2 wild type, *elf3-4*, *phyB-10* and *elf3-4/phyB-10* seedlings under different photoperiodic conditions across time. (B) Hypocotyl growth rate (mm/hr) in the respective genotypes. Highlighted above the plots are the respective photoperiods, long-days in green and short-days in blue. The color boxes highlight the dark phase under the respective photoperiod. All measurements were taken after the photoperiod shift had occurred at ZT72. Note that the seeds germinated later in this experiment than in the experiment depicted in Figure 2, with hypocotyl emergence occurring primarily on day 3.

As ELF3 and phyB were required for photoperiodic memory in seedlings, we investigated whether they contributed to photoperiodic memory in seeds. phyB is functional in seeds (Piskurewicz et al., 2023) and the circadian clock is also active in imbibed seeds (Zhong et al., 1998). The role of ELF3 in seeds remains to be determined. As in WT, the *elf3, phyB* and *elf3/phyB* mutants’ overall growth and the timing of growth showed weak memory of the SD-LD shift (Figure 3A-B). For the LD-SD shift, the *elf3* mutant did not display enhanced growth on the first night (Figure 3B). Instead, the enhanced growth was delayed until the second night. Furthermore, the peak timing of this growth was shifted from late evening to dusk, where ELF3 functions within the evening complex to repress *PIF4/5* expression (Nusinow et al., 2011). As with WT, *elf3* mutants displayed two evenings of accelerated growth following the LD-SD shift before the rate of hypocotyl elongation became more consistent with growth under constant SD (Figure 3B). In the *phyB* mutant, there was no evidence of accelerated growth following from the shift from LD-SD. Instead, growth displayed two smaller peaks at dusk and in the late evening. These two smaller peaks of growth likely compensate for the absence of one predominant period of growth. The *elf3/phyB* double mutant displayed similar phenotypes to the respective single mutants for the LD-SD shift, with both a delay in the adjustment to the new photoperiod and an absence of a major peak of growth (Figure 3B). Thus, phyB and ELF3 seemingly work independently to control the acceleration of growth and adjustment to the new photoperiod, respectively, following the LD-SD shift.

### Circadian clocks adapt to photoperiod shifts quickly in Arabidopsis seedlings

Although our data suggests that seeds and seedlings have a memory of the prior photoperiod, it is unclear how this photoperiodic information is stored. One possible mechanism is through the circadian clock. The circadian clock has a critical role in regulating photoperiodic responses in plants (Nozue et al., 2007; Niwa, Yamashino and Mizuno, 2009; Song et al., 2015; Wang et al., 2024). Furthermore, the circadian clock is active in imbibed seeds (Zhong et al., 1998) and the expression of *PIF4* and *PIF5* display photoperiod-dependent changes in the phase of their expression (Niwa, Yamashino and Mizuno, 2009). Thus, we hypothesized that the circadian clock may function as a repository for photoperiodic information.

To test this possibility, we analyzed the circadian waveform of the *CCR2::LUC* reporter in the WT background. Here, seedlings were entrained for ten days under SD (8/16 light/dark) and then shifted to LD for three days. As a control, seedlings were entrained and then kept under SD or LD for the length of the experiment. For seedlings grown under either constant SD or LD conditions, *CCR2::LUC* was phased to the early evening of each respective photoperiod as reported previously (Strayer et al., 2000; Schultz et al., 2001). In seedlings that underwent the photoperiod shift, the phase of *CCR2::LUC* was initially between the phase of *CCR2::LUC* in either constant SD or constant LD seedlings (Figure 4A). However, by the second day post-shift, the phase of the *CCR2::LUC* reporter became aligned with seedlings that were grown under constant LD. As hypocotyl elongation displays memory of the photoperiod information for at least four days post-shift (Figure 2-3), it is unlikely photoperiodic information stored by the circadian clock can sufficiently explain the developmental responses we have observed.

**Figure 4.**
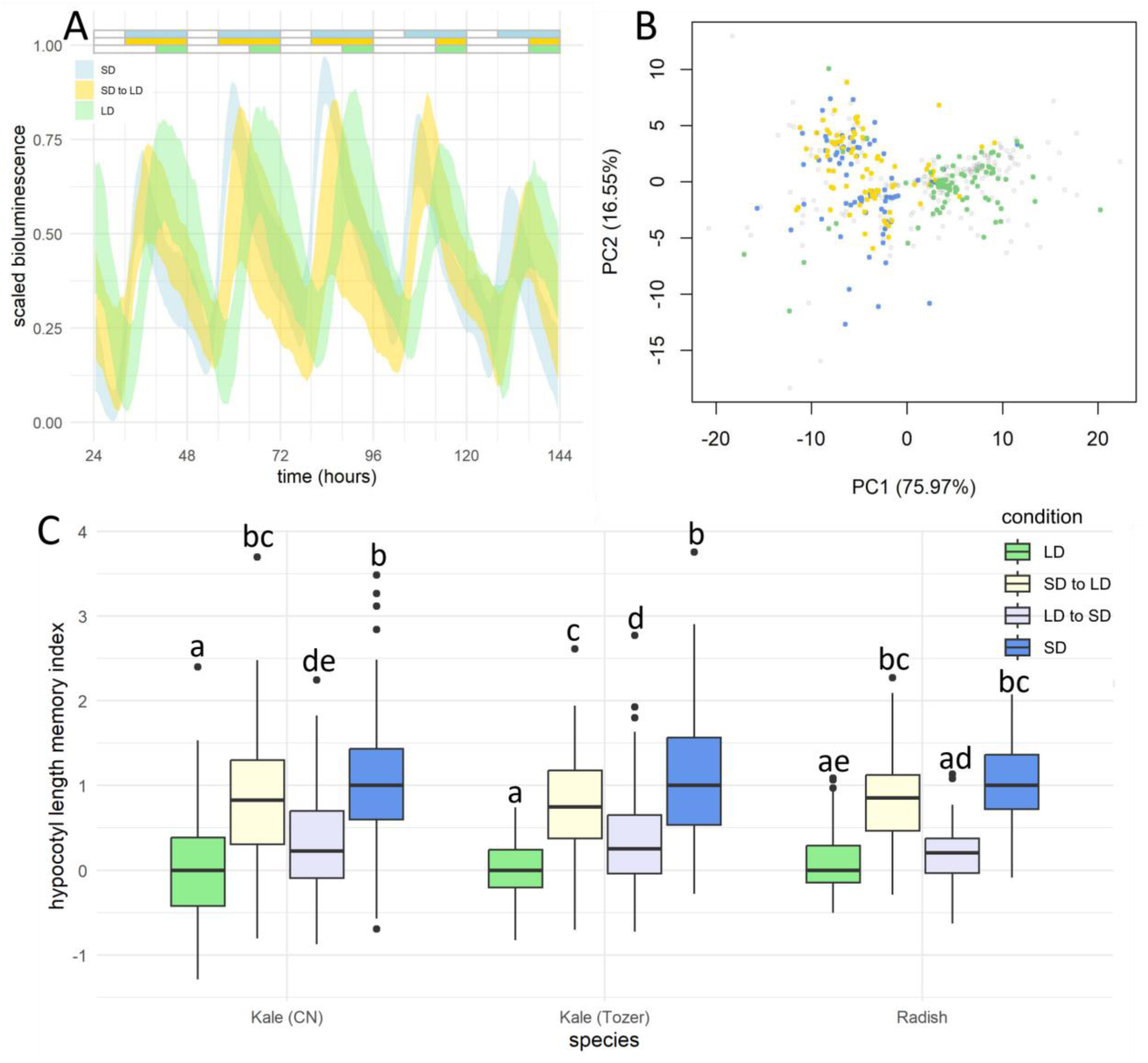
The circadian clock is unlikely to store photoperiodic memory in Arabidopsis. (A) Waveform of *CCR2::LUC* bioluminescence in Ws-2 wild-type seedlings grown under continuous short-day (blue) or long-day (green) or in seedlings shifted from short-day to long-day (yellow). (B) A functional PCA (fPCA) plot, where each point represents a different bioluminescence curve for an individual plant, color-coded by their photoperiod condition that plant was exposed to, using a color-scheme consistent with (A). fPCA is interpreted in the same way as a standard PCA, as a projection of data (in this case, coefficients of a complex B-spline function) to a lower dimensional space. (C) Kale (CN seeds), Kale (Tozer seeds) and radish plants used in microgreen agriculture displayed evidence of photoperiodic memory. Letters signify a significant adjusted p-value of p < 0.01, determined by TukeyHSD.

Although it is unlikely that the circadian clock provides photoperiodic memory, we were interested in understanding the heterogeneity of the response to the SD-LD shift. To investigate this, we visualized the individual responses of each plant that underwent the shift using a functional Principle Component Analysis (PCA). We included three timepoints for each individual plant measured from the day before the shift, the day of the shift, and the day after the shift. Each point represents the shape of the circadian rhythm across the respective day. This plot can be interpreted in the same way as a traditional PCA plot, so days with similarly phased circadian rhythms should group together. We found that there were individual plants that had rhythms that resembled those under LD prior to the SD to LD shift and conversely there were individual plants that had rhythms resembling SD plants two days after the shift (Figure 4B). However, this was a very small proportion of the total plants evaluated. Therefore, the re-adjustment of the circadian phase following the photoperiod shift is robust and uniform. In summary, hypocotyl photoperiodic memory is unlikely to be directly controlled by the circadian clock.

### Other Brassicaceae display strong photoperiodic memory

Next, we investigated whether photoperiodic memory was present in other plants. Here, we focused on microgreens that are grown in vertical farms as the hypocotyl is the primary commercial product (Filatov and Olonin, 2023). Microgreens grown under LD photoperiods have enhanced nutritional quality, but the longer photoperiods are associated with higher operational energy costs. Alongside reducing energy consumption, shorter photoperiods will also induce hypocotyl elongation, increasing the quantity of yield. We hypothesized that photoperiodic memory may enable us to combine the benefit of growth under both photoperiods. To investigate this possibility, we focused on kale and radish crops that are grown as microgreens within vertical farms. As with Arabidopsis, kale and radish displayed evidence of photoperiodic memory (Figure 4C, Supplementary Figure 6). However, all three cultivars that were tested displayed stronger photoperiodic memory than Arabidopsis. Seedlings shifted from SD-LD had a PMI closer to those grown under constant SD, while those shifted from LD-SD had a PMI similar to those grown under constant LD. Moreover, the plants that experienced a SD to LD shift were also less etiolated and had greener coloration, which is a commercially advantageous trait (Supplementary Figure 6). These results suggest that photoperiod shifts may have immediate uses in optimizing growth conditions in vertical farms.

## Discussion

Sensing and responding to photoperiodic changes are essential for plants to adapt to seasonal progression. Hypocotyl elongation is strongly controlled by the photoperiod (Nozue et al., 2007; Niwa, Yamashino and Mizuno, 2009), but it has remained unclear how changes in the photoperiod impacted hypocotyl development. Furthermore, it is unknown if plants store information about prior photoperiodic exposure. Here, we have found that hypocotyl development in different Brassicaceae species displays clear evidence of a photoperiodic memory (Figures 2, 3). Our data directly supports a memory response as there was minimal difference in Arabidopsis hypocotyl length at day 3 between those grown under a SD or LD photoperiod (Figure 1). If there was no memory of the prior photoperiod, then those shifted from SD-LD and vice-versa should closely replicate growth from those under the final photoperiod. However, we did not observe this for any experiment we have carried out. The memory of the initial photoperiod persisted for at least four days in Arabidopsis (Figures 2, 3) and two days in microgreen crops (Figure 4). Different Arabidopsis ecotypes displayed unique sensitivities to the photoperiod shift, but this did not clearly correlate with latitude (Figure 2, Supplementary Table 1). One critical factor in the memory response was the age of the plant when it experienced the photoperiod shift. For instance, shifting germinating Ws-2 seeds from LD-SD resulted in greater hypocotyl elongation relative to those grown under constant SD (Figure 3), a behavior that was not seen when seedlings underwent a similar photoperiod transition (Figure 2). For the SD-LD shift, germinating seeds had a poor memory of the SD-LD compared to seedlings that underwent the same transition (Figures 2, 3). These results highlight how a plant’s response to environmental stimuli needs to be contextualized by the developmental stage.

In the natural environment, seasonal progression results in daylengths changing in the order of minutes, rather than hours. Therefore, it is not clear why plants would need a mechanism for responding to the kind of perturbations we have used in our assay. In natural environments, plants may experience temporary dusk or dawn light disturbances caused by physical obstacles or weather events, which may artificially shorten the day. Plants need a way to distinguish between these kinds of short-term photoperiod fluctuations and the actual photoperiod fluctuations resulting from seasonal progression. In this way, photoperiodic memory may increase robustness to noisy environmental signals. It will be important to understand the effect of smaller photoperiodic shifts and the effect of transient rather than permanent shortening and lengthening of the photoperiod on hypocotyl elongation.

For hypocotyl elongation to be affected by the prior experience of photoperiod exposure, there needs to be an internal mechanism by which information about prior photoperiod exposure is encoded. The circadian clock has an essential role for photoperiodic responses, mediating photoperiod regulation of flowering time (Song et al., 2015) and metabolic growth (Wang et al., 2024). Similarly, the circadian clock has been established as a key regulator of photoperiodic regulation of hypocotyl elongation (Niwa, Yamashino and Mizuno, 2009) and is active in imbibed seeds (Zhong et al., 1998). This led us to hypothesize that the circadian clock may function as a repository for photoperiodic information. However, we found that the circadian clock quickly re-adjusted to the new photoperiod within two days of being transferred and these changes were largely consistent across the population (Figure 4). As photoperiodic memory persisted for at least four days in Arabidopsis seedlings (Figure 2) and seeds (Figure 3), it is unlikely that photoperiodic information is encoded within transcriptional circadian rhythms. Therefore, other mechanisms must exist to store past photoperiodic information to control hypocotyl elongations.

A variety of mechanisms have been described in the literature to explain short-term and long-term memory of environmental stimuli (Murcia et al., 2022; Kambona et al., 2023; Auge et al., 2023). This includes a recent example of how hypocotyl development is controlled by memory of temperature exposure (Murcia et al., 2022). Here, changes in the subcellular localisation of ELF3 function as a device to store a memory of earlier temperature exposure. We have shown that ELF3 is required for photoperiodic memory and is particularly important in regulating the immediate response to both photoperiod shifts (Figures 2, 3). ELF3 can form nuclear bodies at ambient temperatures (Herrero et al., 2012; Anwer et al., 2014) and these are regulated by light, temperature and time (Jung et al., 2020; Ronald, Wilkinson and Davis, 2021; Ronald et al., 2022; Murcia et al., 2022). It will be interesting to investigate whether ELF3 localisation changes under different photoperiods and the role for phyB within this. Additionally, it will be important to further expand our understanding of the genetic pathway responsible for the long-term adaptation to a new photoperiod. Here, phyB seems to have a critical role, with *phyB* mutants displaying reduced growth following either the LD-SD or SD-LD shift (Figure 2). Phytochromes are required for de-etiolation and the switch to photoautotrophic growth in seedlings (Li et al., 2011) and regulate carbon resource allocation in adult plants (Yang et al., 2016). Thus, it will be important to understand the metabolic response of WT and *phyB* mutants to photoperiodic shifts.

### Concluding statement

In summary, our work has uncovered a previously unknown mechanism of photoperiodic memory that controls hypocotyl development. Distinct pathways seem to operate to control the response to different directions of photoperiodic shifts, and the impact of the memory is dependent on the developmental age of the plant. The genetic pathway that regulates photoperiodic memory is unclear, but it seems to occur independently of transcriptional circadian rhythms. Photoperiodic memory may have immediate direct agricultural implications by allowing the optimization of nutritional quality, yield, and operational costs of vertical farms. Future work can explore whether photoperiodic memory persists in non-Brassicaceae species to understand the wider potential benefits of photoperiodic memory in other commercial applications.

## Figures

**Supplementary Figure 1 –** Representative scans of seedlings from wild-type Arabidopsis ecotypes used in this work. Images are from day 5, two days after the photoperiod shift. Scale bar is 5 mm.

**Supplementary Figure 2 –** Hypocotyl length (mm) of wild-type ecotypes sorted by total light exposure (hours) on day 5. Dark blue is constant short-day, purple are seedlings shifted from long-day-short-day, yellow is seedlings shifted from long-day to short-day and green is constant long-day.

**Supplementary Figure 3 -** Hypocotyl length (mm) of wild-type ecotypes sorted by total light exposure (hours) on day 7. Dark blue is constant short-day, purple are seedlings shifted from long-day to short-day, yellow is seedlings shifted from long-day to short-day and green is constant long-day.

**Supplementary Figure 4 –** Representative scans of seedlings from wild-type Arabidopsis Ws-2, *elf3-4*, *phyB-10* and *elf3-4/phyB-10.* Images are from day 5, the first measurement point. Scale bar is 5 mm.

**Supplementary Figure 5 -** Hypocotyl length (mm) of wild-type Ws-2, *elf3-4*, *phyB-10* and *elf3-4/phyB-10* sorted by total light exposure (hours) on (A) day 5 and (B) day 7. Dark blue is constant short-day, purple are seedlings shifted from long-day-short-day, yellow is seedlings shifted from long-day to short-day and green is constant long-day.

**Supplementary Figure 6 -** Representative scans of microgreen seedlings. Microgreens are approximately 11 days old at the end of the experiment Scale bar is 15 mm.

**Supplementary File –** Raw data and statistical tests associated with the figures of this manuscript.

## Acknowledgements

We would like to thank Maarten Koornneef for providing the Tnz-1 seeds. We would also like to acknowledge A. L. Tozer Limited for providing us with radish and kale seeds and the York Biology horticulture staff for their assistance and support with this work.

---

**Supplementary Table 1.** Arabidopsis ecotypes used in this work.

| Ecotype | Latitude | Longitude | Country |
| --- | --- | --- | --- |
| Wassilewskija-2 (Ws-2) | 52.3 | 30.0 | Belarus |
| Tanzania (Tnz) | -2.9 | 36.2 | Tanzania |
| Bay-0 | 49.0 | 11.0 | Germany |
| Shakdara (Sha) | 38.4 | 68.5 | Tajikistan |
| Cape Verde Island (Cvi) | 15.1 | -23.6 | Cape Verde Island |
| Sapporo (Sap) | 43.1 | 141.3 | Japan |

**Supplementary Table 2.** Microgreen cultivars used in this work.

| Microgreen | Species | Variety |
| --- | --- | --- |
| Kale (Tozer Seeds) | *Brassica oleracea* | Dwarf Green Curled |
| Kale (CN Seeds) | *Brassica oleracea* | Dwarf Blue |
| Radish | *Raphanus raphanistrum* | Sangrin |

