## Supplemental Materials for "Hypocotyl Development in Arabidopsis and other Brassicaceae Displays Evidence of Photoperiodic Memory"

### Slide 1
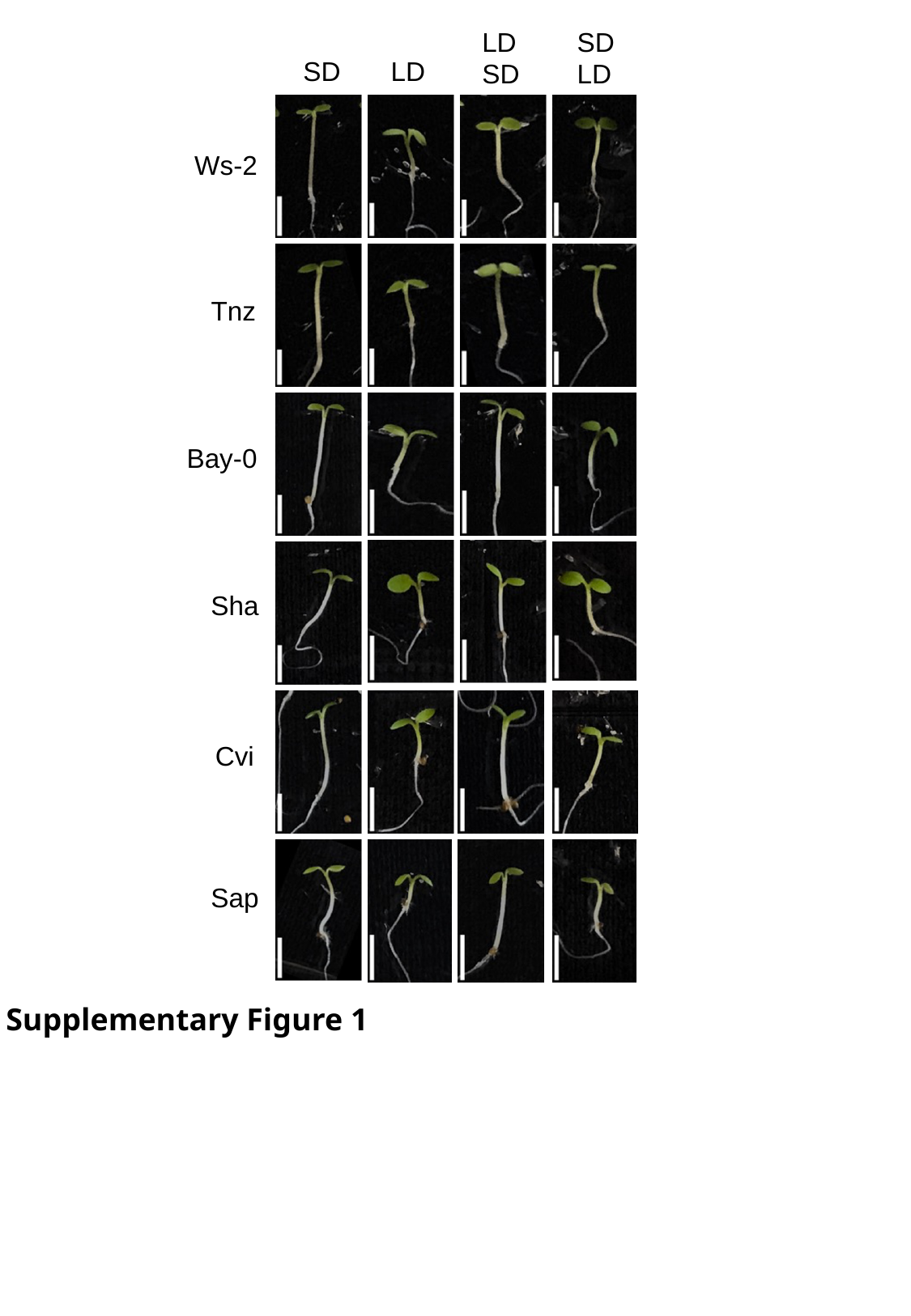

SD
LD
LD
SD
SD
LD
Ws-2
Tnz
Bay-0
Sha
Cvi
Sap
Supplementary Figure 1

### Slide 2
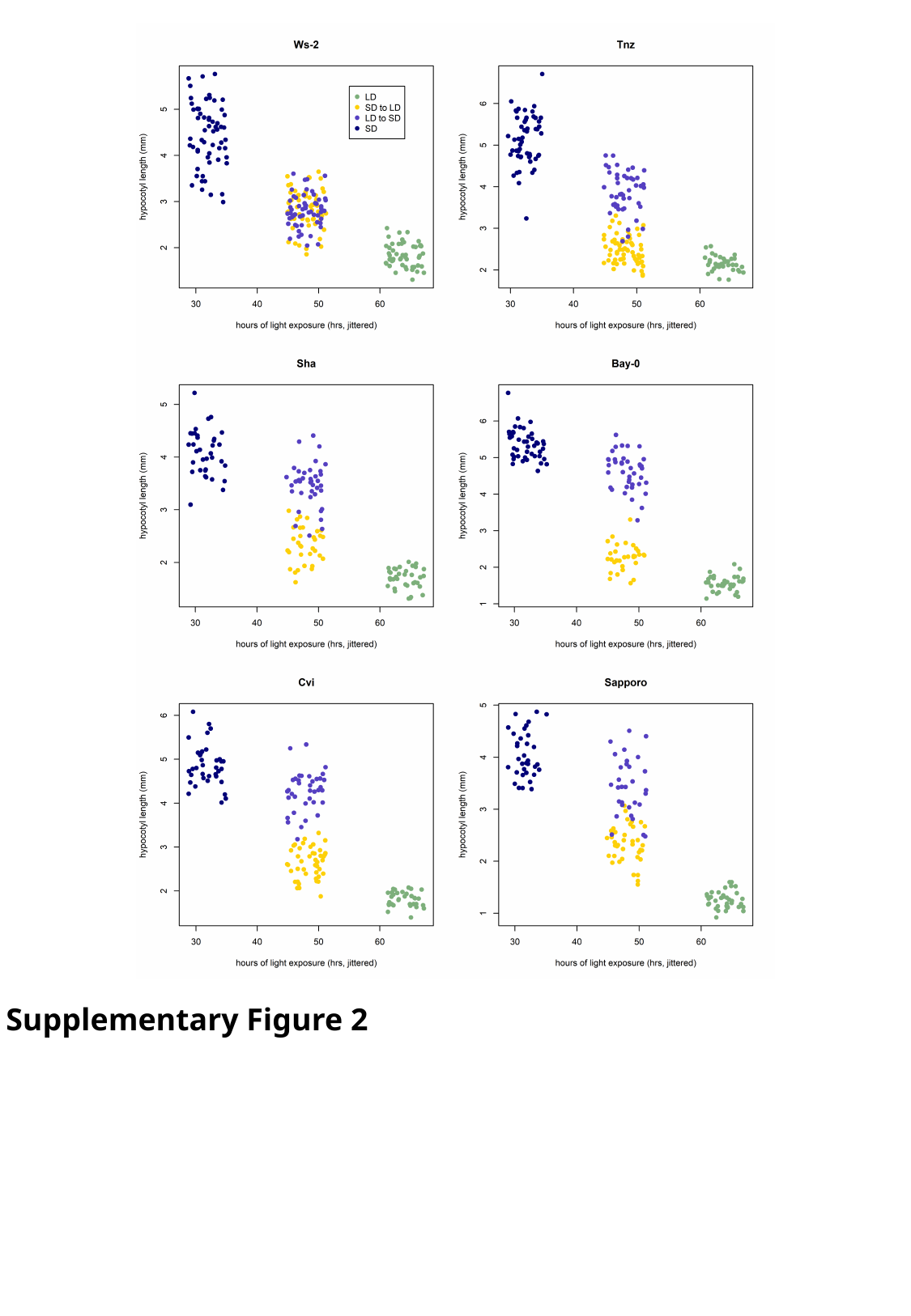

Supplementary Figure 2

### Slide 3
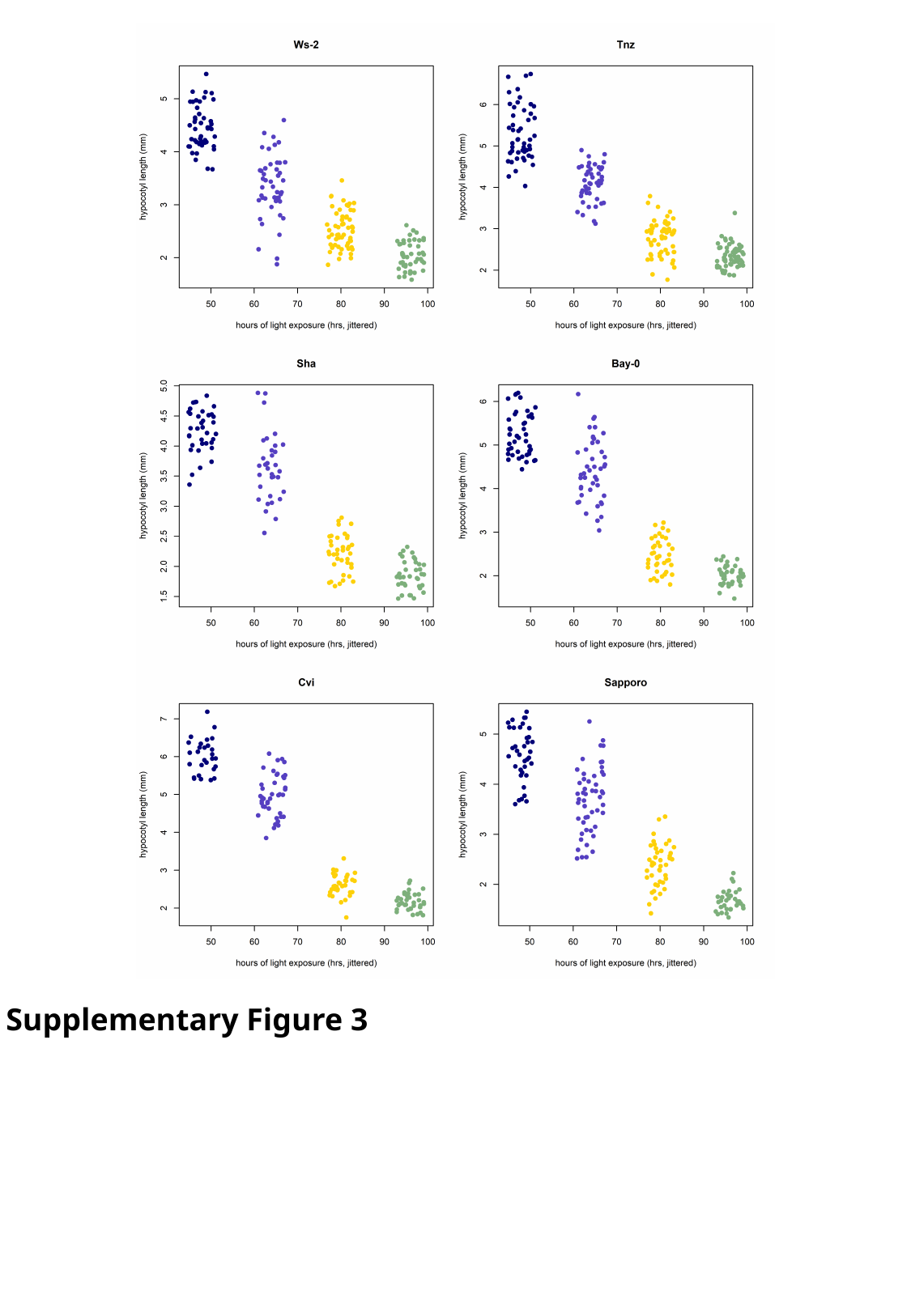

Supplementary Figure 3

### Slide 4
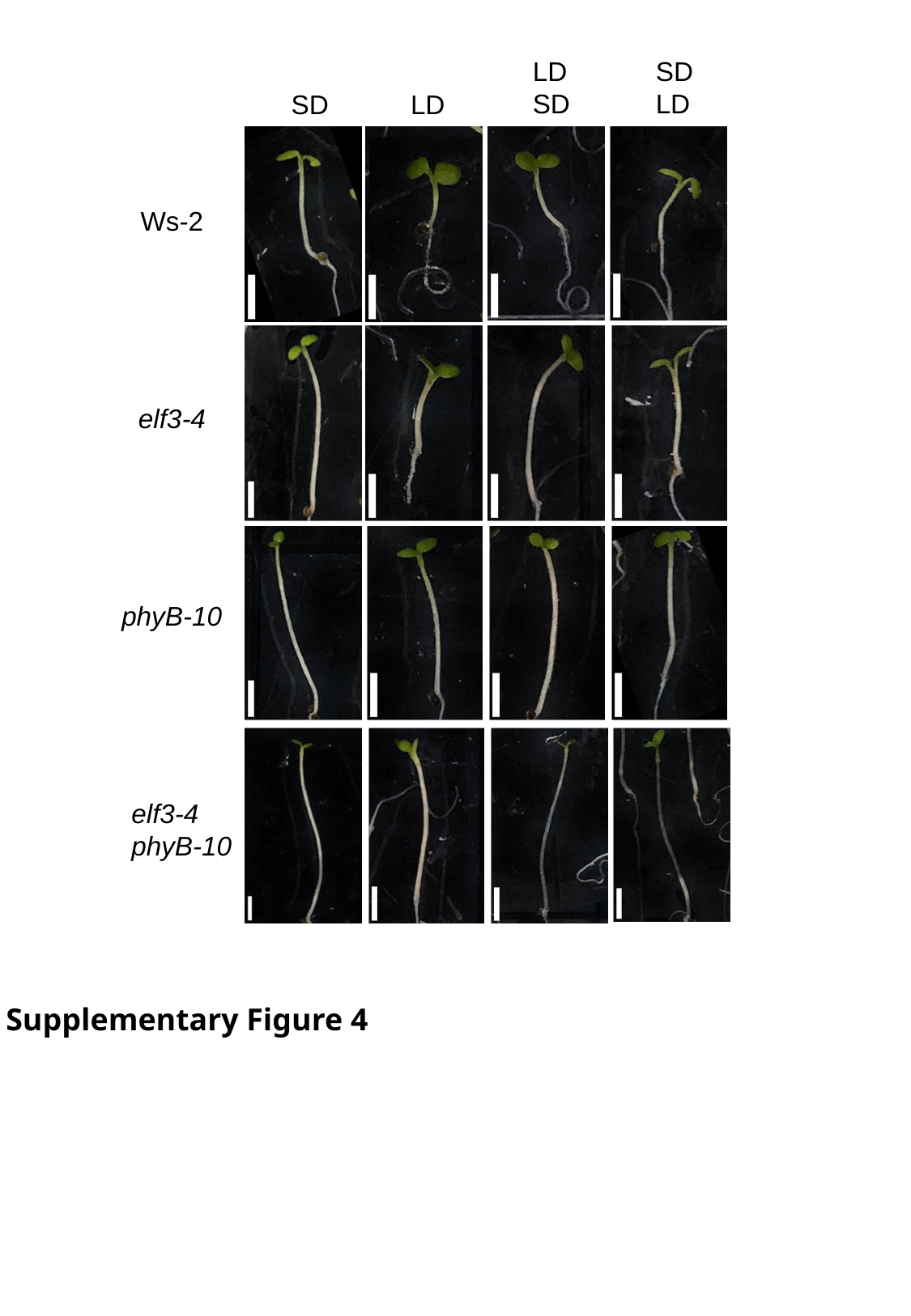

LD
SD
SD
LD
LD
SD
Ws-2
elf3-4
phyB-10
elf3-4
phyB-10
Supplementary Figure 4

### Slide 5
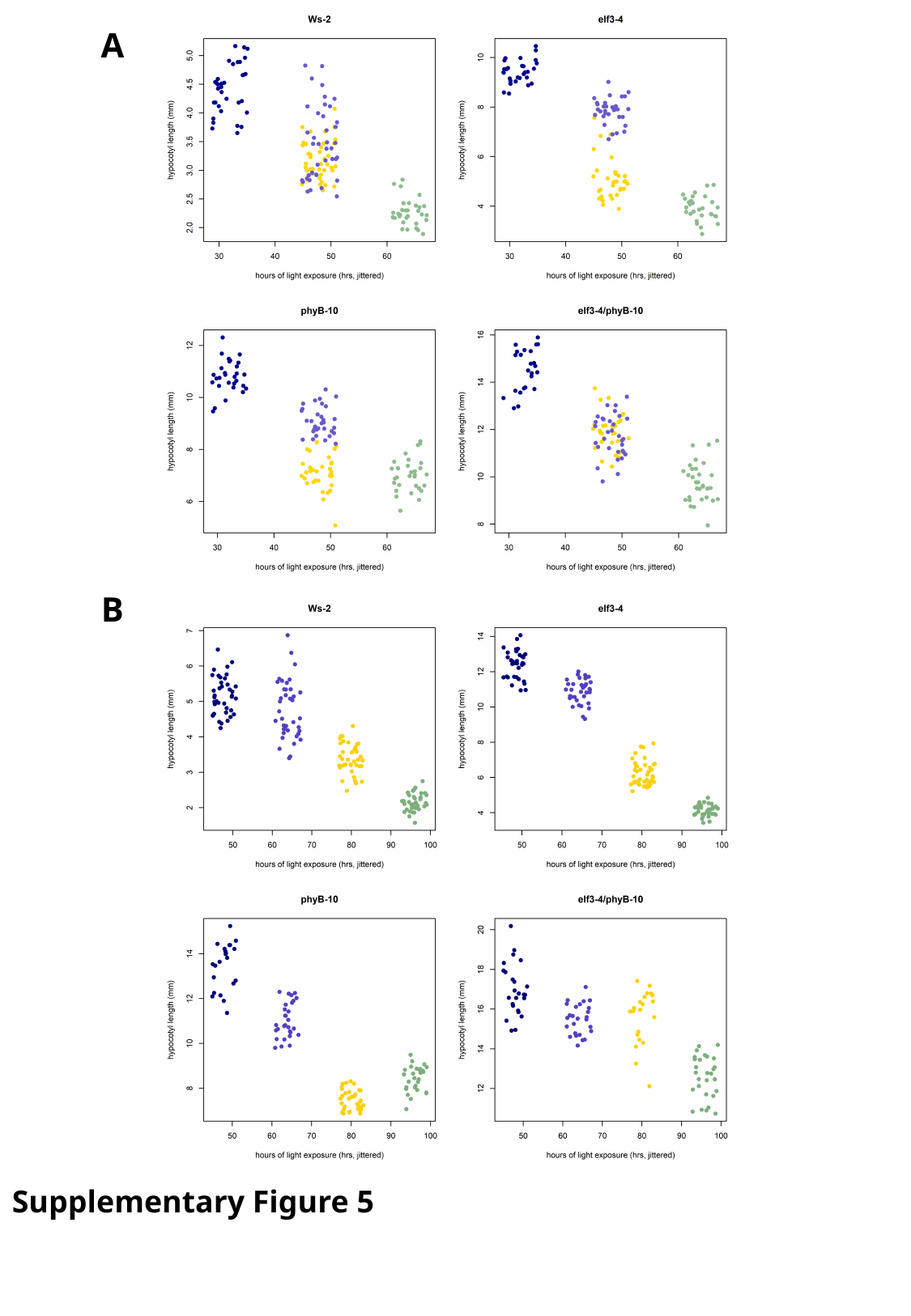

A
B
Supplementary Figure 5

### Slide 6
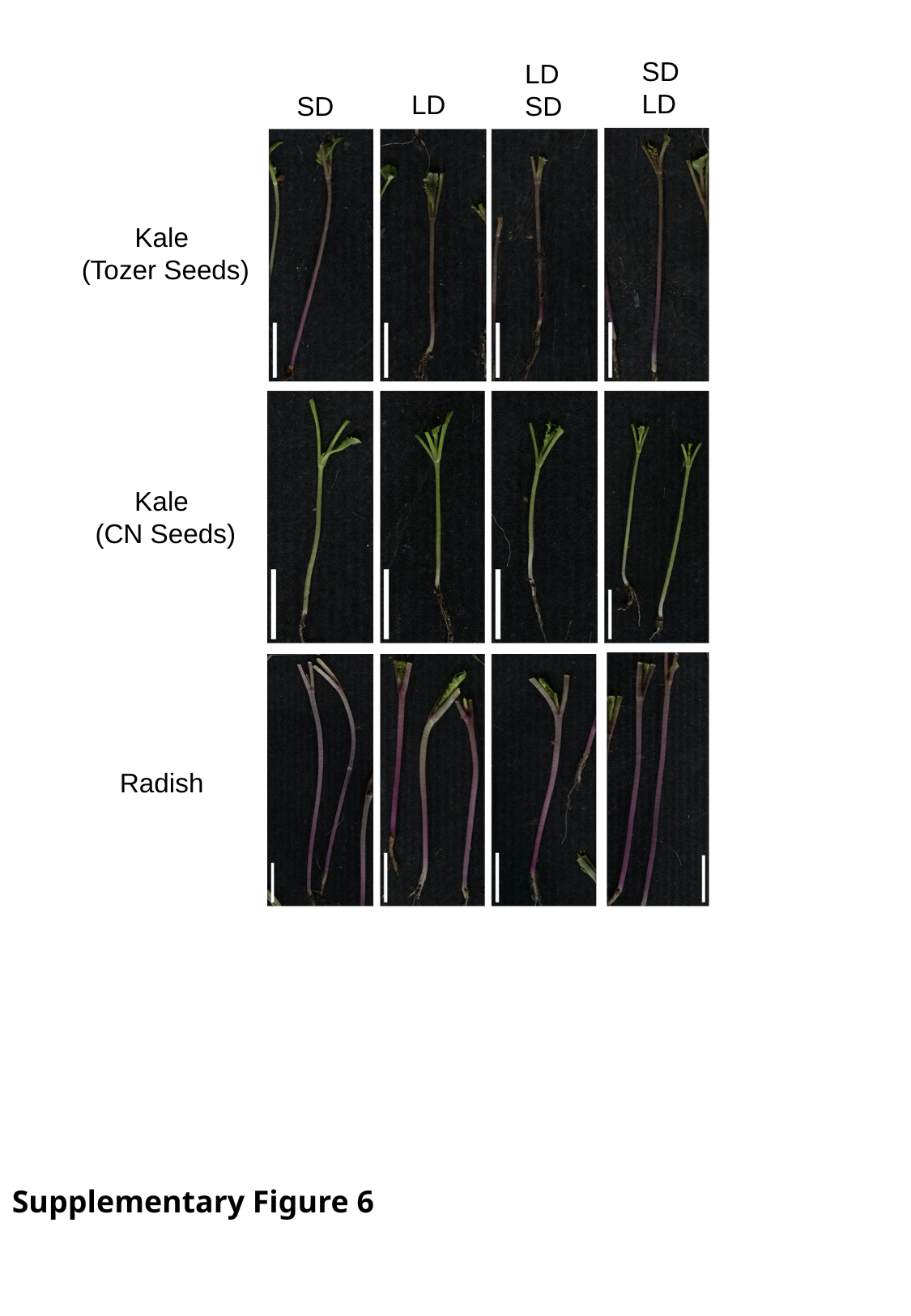

SD
LD
LD
SD
LD
SD
Kale
(Tozer Seeds)
Kale
(CN Seeds)
Radish
Supplementary Figure 6
